# Velocity and Diameter Measurements of Penetrating Arteries by Model Based Analysis of Complex Difference Images in Phase Contrast MRI

**DOI:** 10.1101/293308

**Authors:** Xiaopeng Zong, Weili Lin

## Abstract

Pathological changes of penetrating arteries (PAs) within deep white matter (WM) may be an important contributing factor of cerebral small vessel disease (SVD). Quantitative characterization of the PAs is important for further illuminating their roles in SVD but remains challenging due to their sub-voxel sizes. We propose a quantitative MRI approach for measuring the diameters and flow velocities of PAs based on model based analysis of complex difference images in phase contrast MRI. The complex difference image of each PA is fitted by a model image calculated by taking into account the partial volume effect and signal enhancement due to in flow effects to obtain velocity 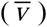, diameter (*D*), and volume flow rate (VFR) of the PAs. Simulation, phantom, and in vivo studies were carried out to evaluate the accuracy and measurement errors of the proposed method. Our results suggest that PAs with velocities ≥ 0.8 cm/s can be accurately measured with 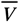, *D*, and VFR errors of 0.28 cm/s, 20 μm, and 0.024 mm^3^/s, respectively, although the mean lumen area occupies only 18% of the acquired pixel area. The PAs have a 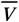 distribution peak at ~1.2 cm/s and diameters distribution mostly in the range of 88 – 200 μm Quantitative measurements of PAs with the MBAC method may serve as an invaluable tool for illuminating the role of PAs in the aetiopathogenesis of cerebral SVD.

## Introduction

Pathological changes of penetrating arteries (PAs) within deep white matter (WM) of the centrum semi-ovale (CSO) is an important contributing factor of cerebral small vessel disease (SVD).(Fisher 1968, Jorgensen, Shaaban et al. 2018) The main effects of the pathological changes include narrowing or occlusion of the vessels leading to focal ischemia, and leakage of blood brain barrier leading to accumulation of toxic blood components in surrounding tissue.(Fisher 1968, Wardlaw, Dennis et al. 2001, Montagne, Nikolakopoulou et al. 2018) Measuring the morphological and functional properties of the PAs can help differentiate the underlying pathology of PAs,(Wardlaw, Sandercock et al. 2003, Bernbaum, Menon et al. 2015) which may in turn lead to the development of effective treatment strategies for improving PA function.

Phase contrast (PC) MRI is an established method for measuring blood flow in large arteries such as internal carotid and basilar arteries.(ten Dam, van den Heuvel et al. 2007, van der Veen, Muller et al. 2015) However, most PAs have diameters of 40 – 500 μm.(Pesce and Carli 1988) Therefore, there exists strong partial volume effects in the images with currently achievable MRI spatial resolutions. Different image analysis strategies have been proposed to account for partial volume effects in flow quantification. Tang et al. proposed to correct the volume flow rate in voxels near the boundary of the lumen by the ratio of image intensity in such voxels to that of voxels fully occupied by blood.(Tang, Blatter et al. 1995) However, this method is not applicable to our case since no voxel fully occupied by blood exist for the PAs. Hamilton proposed to fit a circle to the complex images acquired with and without the velocity encoding gradients, respectively.(Hamilton 1994) However, this method does not consider the Gibbs ringing effects due to the finite extent of k-space sampling. Lagerstrand et al. proposed to correct the partial volume effects by a calibration curve obtained from a separate measurement on a flow phantom with similar relaxation and flow properties.(Lagerstrand, Lehmann et al. 2002) Due to the requirement of a calibration curve, this method cannot be readily applied to in vivo studies with unknown flow rates. Hoogeveen et al. proposed a model-based analysis of the phase images (MBAP),(Hoogeveen, Bakker et al. 1999) where the model image accounts for blurring due to limited spatial resolution and flow-induced signal enhancement. However, the accuracy of the MBAP approach for sub-pixel vessels was not systematically evaluated, and the error for VFR already reached ~45% when the pixel dimension was only slightly greater than (1.1 times) the vessel diameter. Furthermore, a two-pool model was assumed in calculating the model phase images, which may not be applicable for the PAs due to the presence of surrounding perivascular spaces and white matter with very different MR properties.(Zong, Park et al. 2016)

In this study, we aim to develop a new approach for measuring diameter and flow velocity of PAs by model-based analysis of complex difference images (MBAC). Complex difference (CD) eliminates the static tissue contributions in the images. Therefore, our approach is applicable even when there are multiple static tissue pools. Furthermore, compared to the MBAP method, the MBAC approach utilizes both the phase and magnitude changes induced by flow for improving the precision of fitted parameters. We present detailed simulation, phantom, and in vivo studies for evaluating the precision and accuracy of the new technique.

## Methods and Materials

### Theory

Assuming the vessels are oriented perpendicular to the slice, the MR signal in a PC MRI scan can be expressed as:

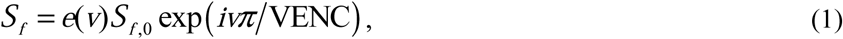

where *S*_*f,0*_ is the steady state signal without inflow effect, *v* the flow velocity, *e*(*v*) the flow induced signal enhancement factor. In PAs, the Reynold number (R) is ~0.3, which is much less than the critical values for turbulent flow,(Avila, Moxey et al. 2011) assuming a blood kinetic viscosity of ~3 × 10^−6^ m^2^/s (https://physics.info/viscosity/), velocity of 1 cm/s, and diameter (D) of 100 μm. Therefore, the blood velocity pattern can be approximated by a laminar velocity distribution: 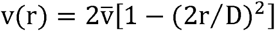, where 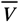 is the mean velocity. Errors caused by deviation from the laminar pattern will be evaluated by simulations. Assuming k-space sampling on a Cartesian grid, the acquired image is a discrete sampling of the total MR signal spatial distribution after convolution with sinc shaped point spread functions:

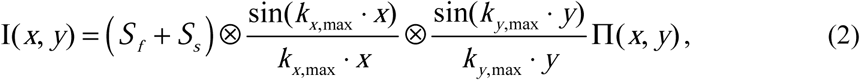

where *k*_x,max_ and *k*_y,max_ are the largest sampled k-space coordinates, П(*x*, *y*) takes nonzero unit values at an equally spaced two dimensional grid, and *S*_*s*_ the steady state signal of the static spins. The CD of *I* (*x*, *y*) with and without velocity encoding gradients removes the static spin contributions and is given by:

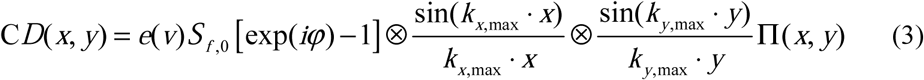

Experimentally, CD can be calculated either directly as

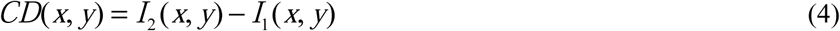

where *I*_1_ and *I*_2_ are the complex images with and without velocity encoding gradients, or under certain conditions as

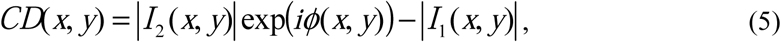

where *φ* is the phase difference between images with and without flow encoding and is calculated as

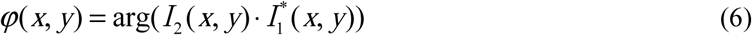

For phase array coils, the product in Eq. (6) is calculated for each coil and then the average value is used for calculating the phase difference.(Bernstein, Grgic et al. 1994) Equation (5) is more convenient to calculate than Eq. (4) for phased array coils since there is no need to reconstruct phase images for *I*_1_ and *I*_2_. These two equations are equivalent when *I*_1_ has a zero phase (after removing constant offset and slow spatial variations which can be induced by eddy-current). However, Eq. (5) is no longer valid if the phase oscillates due to Gibbs ringing effect. The phase will change by 180° when negative Gibbs ringing signal from the vessel exceeds the signal of surrounding tissue.

The flow enhancement factor *e*(*v*) is needed for calculating the model images, which depends on the slice selection (SS) profile of the RF pulses. For any realistic RF pulse, the flip angle (θ) varies along the slice direction. Therefore, *e*(*v*) can only be obtained numerically. In our study, a five lobe sinc pulse apodized with a Hamming window was employed for excitation. The SS profile (i.e. θ versus position (*z*) in slice direction) of the pulse was calculated via Bloch simulation. The resulting profile of the RF pulse used in our experiment is shown in Figure 1(A) together with the boxcar profile of an ideal pulse. The MR signal at position *z* is given by

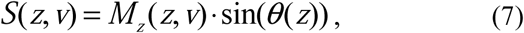

where *M*_*z*_ is the position and velocity dependent longitudinal magnetization which can be calculated iteratively from the following equation:

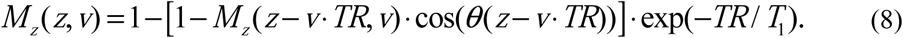

As *θ*(z) is essentially zero outside [−1.5*t*, 1.5*t*], where *t* is the slice thickness, *M*_*z*_(*z*,*v*) can be calculated iteratively by setting *M*_*z*_(*z*, *v*) = 1 for *z* ≤−1.5*t*. *e*(*v*) is calculated as the ratio of mean S(*z*,*v*) over *z* ∈ [−1.5*t*, 1.5*t*] to the mean S(*z*, 0) over the same *z* range, i.e.

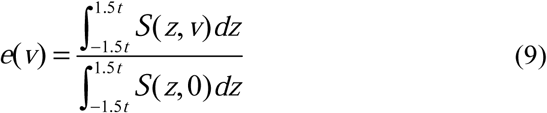

In Figure 1(B), *e*(*v*) is plotted for the apodized sinc and ideal pulses, assuming blood T_1_ = 2.6 s and TR = 26 ms. A strong dependence of *e*(*v*) on SS profile is observed in Figure 1B, demonstrating the importance of employing correct SS profile in the calculation.

**Figure 1.**
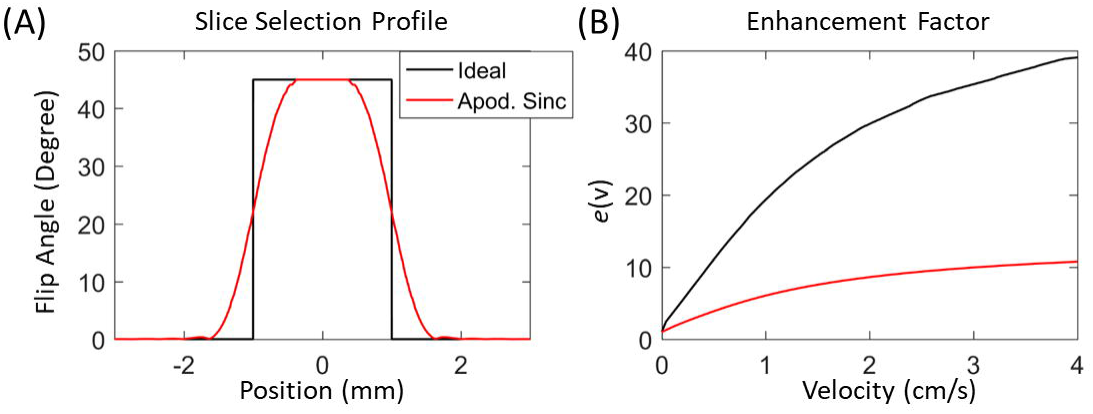
(A): Slice selection profiles of the apodized sinc pulse. An ideal pulse with boxcar profile is also shown for comparison. (B): Flow enhancement factors versus flow velocity calculated for the two profiles for flow direction perpendicular to the imaging slice. The blood T_1_ is assumed to be 2.6 s and TR = 26 ms.

Perpendicular vessels were assumed in the equations (1), (8), and (9). When the vessels deviate from the slice normal direction, Eq. (1) needs to be replaced by

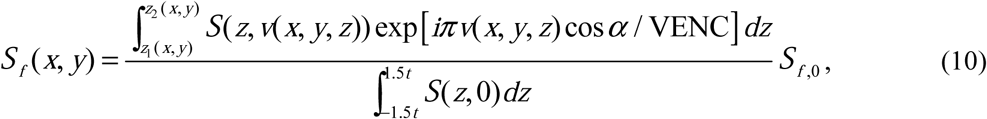

where *v* is now *z*-dependent due to the vessel tilt, α is the angle between the vessel and normal directions, and the integration range [z_1_, z_2_] denotes the intravascular segment of the integration path [−1.5t,1.5t]. Equation (8) also needs to be modified to account for the vessel tilt as follows:

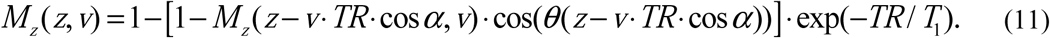

The steady state arterial blood signal (*S*_*f,0*_) cannot be measured directly but can be calculated from the steady state signal of the surrounding WM (*S*_*wm*_) via the following equation:

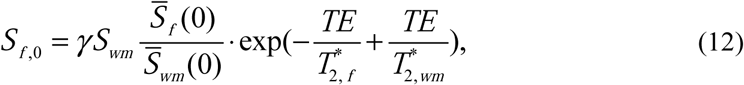

where γ is the blood-WM water partition coefficient, 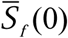 and 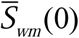 are mean *S*(*z*,*0*) over *z*∈[−1.5*t*, 1.5*t*] for the arterial blood and WM spins, respectively, and 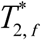 and 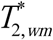 are their 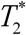 values.

The MBAC method estimates 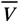, *D*, and volume flow rate (VFR; which is 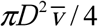) by fitting the experimental CD image calculated from Eqs. (4) or (5) with the model CD image calculated from Eq (3). Alternatively, the phase image ϕ (*x*, *y*) in Eq. (6) can also be fitted with a model image calculated with Eqs. (2) and (6), as first demonstrated by Hoogeveen et al(Hoogeveen, Bakker et al. 1999) and will be denoted as MBAP in our paper. Simulations will be carried out to compare the noise-induced estimation errors of the two approaches. Furthermore, note that perpendicular vessels are assumed in calculating the model CD and phase images. Simulation will be also carried out to evaluate errors caused by deviation of the vessels from the perpendicular direction in the MBAC method.

### Simulation

We carried out simulations to evaluate the errors of fitted 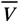, *D*, and VRF due to noise, deviation of vessel orientation from slice normal direction, and non-laminar velocity patterns. True *D* (*D*_true_) was between 80 and 200 um and true 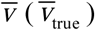 was between 0.4 cm/s and 2 cm/s. For each *D*_true_ and 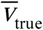 combination, two simulated image were calculated with VENC = 4 cm/s and ∞ (i.e. velocity encoding gradient off), respectively. The assumed MR parameters closely matched with *in vivo* condition: *k*_*x*,max_ = *k*_*y*,max_ = 10.05 mm^-1^ corresponding to acquired pixel size 0.3125×0.3125 mm*2*, *T*_1,*f*_ = 2.6 s for blood at 7 T (Rooney, Johnson et al. 2007)), T_1,*s*_ = 1.2 s for white matter (Rooney, Johnson et al. 2007), 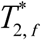 and 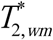 equal to 29 ms and 24 ms, respectively, *TE* = 15.7 ms, *TR* = 26 ms, γ=1.05, *θ*(0) = 45°, and *θ*(z) matching the experimental RF slice selection profile. The 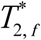 and 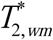 values were measured at internal carotid arteries (ICA) and WM ROIs in a single young healthy subject (age 21, male) using a multi-echo gradient echo sequence and standard data fitting procedure. The convolution in Eq. (2) was implemented as finite element calculation with element width equal to 1/64 of the acquired pixel width. The five central lobes of the sinc functions were taken to speed up the calculation, as the errors due to truncation were found to be negligible.(Hoogeveen, Bakker et al. 1999) The convoluted high resolution images were downsampled by a factor of 32 to produce the final simulated images with pixel size of 0.1563×0.1563 mm^2^ and matrix size of 11×11.

To evaluate errors due to noise, simulated images were calculated assuming perpendicular vessels orientation and laminar flow pattern. Complex noises were added to the simulated images such that the resulting SNR of the WM was 45, consistent with *in vivo* images. Based on the resulting images, CD and phase images were calculated from Eqs. (4) (for MBAC) and (6) (for MBAP), respectively, and fitted with model images with 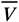, *D*, and vessel position as free parameters. The model images were calculated using the same parameters as for the simulated images before noise addition, except for the fitting parameters. The fitting procedure was repeated 100 times for each *D*_true_ and 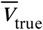 combination with new noise generated for each repetition to calculate the mean and standard deviations (SD) of the fitted 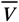, *D* and VFR.

In addition to performing MBAC and MBAP model fitting, the above simulated images were also used for calculating apparent 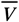, *D*, and VFR in order to demonstrate the large of partial volume effect on the vessel parameter estimation. In order to calculate the apparent parameters, PA ROIs were first defined based on the simulated phase difference and magnitude images. Specifically, possible PA pixels were first defined in a 4×4 square at the image center that have the phase difference or magnitude 1.96σ above the mean background values, where σ is the expected standard deviation of the phase difference or magnitude due to the added noise. Then the possible PA pixels were grouped into neighboring clusters and the largest cluster in the phase difference images that overlapped with a cluster in the magnitude image were defined as the PA ROI in that repetition. Apparent 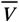 was calculated as the mean velocity of all pixels in the ROI, based on the apparent phase difference. Apparent *D* was calculated as 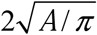, where *A* is the total area of the ROI.

To evaluate errors caused by oblique vessels, simulated images were calculated assuming nonzero angles between the vessel and the slice normal direction, and a laminar flow pattern. Then, the same noise addition and fitting procedure as above was carried out and repeated for 100 times under each tilt angle, *D*_true_ and 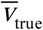 combination. The model images were calculated using the same parameters as for the simulated images before noise addition, except for the fitting parameters and fixing the tilt angle to zero. The fitted 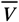, *D* and VFR values from the 100 repetitions were averaged across the repetitions and then compared with their true values.

To evaluate errors caused by deviation of the velocity distribution from laminar flow pattern, simulated images were also calculated assuming a plug flow pattern with uniform velocity distribution and perpendicular vessels. Then, the same noise addition and fitting procedure as above were carried out for 100 times for each *D*_true_ and 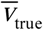 combination. The model images were calculated using the same parameters as for the simulated images before noise addition, except for the fitting parameters and assuming a laminar flow pattern. The fitted 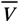, *D* and VFR values were averaged across the repetitions for comparison with their true values.

The model fitting was carried out with the lsqnonlin function in MATLAB version R2016b (MathWorks, Natick, MA, USA). Only pixels within a circular ROI centered at field of view (FOV) center and with a diameter of 9 pixels were included in the fitting. The fitting function returns when the final step size was less than 10^−6^, with position and *D* parameters in mm and 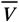 in cm/s. To ensure satisfying the convergence criteria for all repetitions, the “FunctionTolerance” and “OptimalityTolerance” options were set to 0 and the “MaxFunctionEvaluations” and “MaxIterations” options were set to infinity. In almost all cases, the function returns after less than 100 iterations. The initial values for 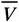, *D* were set to 90% of their true values while the initial vessel position matches the true vessel location.

### MRI experiments

All experiments were performed on a Siemens 7T MRI human scanner (Siemens Healthineers, Erlangen, Germany) using an eight-channel transmit and thirty two-channel receive coil (Nova Medical, Wilmington, MA, USA). Images were acquired with a single slice 2D PC MRI sequence. The velocity encoding gradients were turned on and off alternatively in subsequent TRs to acquired images with and without flow-induced phase shifts, respectively. B_1_ field calibration was carried out with a vendor-supplied presaturation-based B_1_ mapping sequence.

#### Phantom Study

##### Phantom preparation

A flow phantom was constructed to evaluate the accuracy of the MBAC method. The phantom consisted of a Polyethylene PE-10 tube (ID = 0.28 mm; OD = 0.61 mm) penetrating horizontally through a cylindrical plastic cup (ID 5 cm; H = 6.5 cm). The tube was horizontal and parallel to the base of the cup. Both the cup and tube were filled with tap water. The water flow within the PE-tube was driven by a 5 ml syringe connected to the end of the tube. The syringe was pushed by a syringe pump at six constant rates between 0.554 ml/h and 3.33 ml/h to achieve mean flow velocities of 0.25 cm/s to 1.5 cm/s in step of 0.25 cm/s.

##### MRI Parameters

The PC MRI sequence parameters were: VENC = 4 cm/s; FOV = 153×124 mm^2^; matrix = 256×208; thickness = 2 mm; in-plane resolution = 0.6×0.6 mm^2^; flip angle: 15°, 25° or 35°; TR/TE = 26.0/15.7 ms; Number of averages (NA) = 16; slice perpendicular to the tube. Water T_1_ needed for *e*(*v*) calculation was measured with a variable TR turbo-spin echo sequence with 9 TR values in the range of 0.28 s – 18 s.(Conturo, Beth et al. 1987)

##### Data analysis

Images were reconstructed offline to a pixel size of 0.3×0.3 mm^2^, i.e. half the acquired pixel size by zero padding in k-space before inverse Fourier transform. The magnitude images from each coil were combined with root mean square. The phase images were calculated from the complex images of all coils (denoted as *I*_*j*_) as

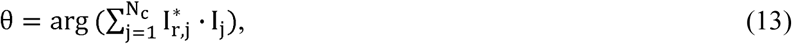

where *N*_*c*_ is the total number of channels and *I*_*r,j*_ is the image without flow encoding gradient for coil j after aligning the mean phases of all channels in a ring ROI centered on the tube. The ring ROI had inner and outer radii of 1.2 mm and 2.7 mm, respectively, and covered the region of stationary water.

The phases within all pixels in the ring ROI were fitted with a second order polynomial of the pixel position to estimate the background phase spatial variations. The estimate background phase was then subtracted from the phase images. The resulting phase images were then used for calculating the CD image with Eq. (4). Eq. (5) was not used as there was no signal within the vessel wall, so the phase within the vessel wall may change by 180°. The CD of pixels within a circular ROI with radius of 1.2 mm were fitted with a model CD image to obtain 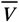 and *D*. The model images were calculated assuming perpendicular vessels and a laminar flow pattern. In addition to 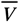 and *D*, other fitting parameters include the vessel center coordinates. The details for the model fitting routine are the same as described in the simulation. As the flowing and stationary spins are both tap water, *S*_*f,0*_ is equal to the mean signal in the ring ROI defined above. The water T_1_ was obtained by fitting the standard longitudinal magnetization recovery equation to mean signal within a circular ROI of 1-cm diameter outside the tube versus TR curve.

Apparent 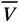, *D* and VFR were also calculated for the flow phantom, in the same way as described as for the simulated images, to demonstrate the strong partial volume effects. However, because of the lack of signal from the tube wall which results in low signal in the magnitude images at the vessel, the vessel ROIs were defined only based on the phase difference images. Specifically, vessel pixels were first defined as those within the above 1.2 mm circular ROI that has phase difference 1.96 times the standard deviation above the mean background values. The standard deviation and mean background values were calculated from the pixels in the ring ROI described above.

#### In vivo experiments

*In vivo* experiments were carried out to evaluate the reproducibility of 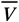, D, and VFR measurements in PAs in CSO and to evaluate the accuracy of MBAC results in ICA where true 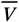, D, and VFR can be obtained.

##### Subjects

Ten healthy volunteers participated in the study; six (ages: 21–41; 4 male) for imaging PA and the other four (ages: 26 – 43; all female) for imaging ICA. The study was approved by the institutional review board at University of North Carolina at Chapel Hill. Written informed consent was obtained from each subject before the experiments.

##### Data acquisition

The sequence parameters for PA imaging were: VENC = 4 cm/s; FOV = 200×162.5 mm^2^; matrix: 640×520; thickness = 2 mm; in-plane resolution: 0.3125×0.3125 mm^2^; GRAPPA factor = 2; TR = 25.5 – 30.0 ms; TE = 15.7 ms; NA: 15-20; axial slice ~1.5 cm above the corpus callosum; flip angle: 30° (n=1), 45° (n=4) or 60° (n=1). Large flip angles (i.e. > Ernst angle) were chosen to increase in-flow signal enhancement and suppress white matter signal. The three angles were found to produce similar vessel visibility and thus their results will be combined in the group analysis. The slice thickness of 2 mm was chosen to be less than the lengths of most PAs. Based on the length distribution of perivascular spaces surrounding PAs, the majority (75%) of PAs should have length ≥ 4 mm (Zong, Park et al. 2016). Scans were repeated twice during the same session for evaluating the reproducibility.

Segmentation of CSO in the images can help exclude false PAs located on the brain surface. To facilitate CSO segmentation, a T_1_-weighted 2D inversion recovery gradient echo sequence was performed at the same slice location as the above PC MRI sequence. MRI sequence parameters of the inversion recovery sequence were: TI/TR/TE = 1100/2200/3.72 ms; matrix: 200×184; in-plane resolution: 1×0.88 mm^2^; lines per TR: 23.

The sequence parameters for ICA imaging were: VENC = 90 cm/s; FOV = 200×162.5 mm^2^; matrix size: 448×364; thickness = 2 mm; in-plane resolution: 0.446×0.446 mm^2^; GRAPPA factor = 2; flip angle = 45°; TR/TE = 30.0/5.7 ms; NA = 18; axial slice cutting through a vertical segment of the ICA as visualized on pilot 3D MPRAGE images.

##### Data Analysis

*in vivo* images for PAs were reconstructed offline to pixel sizes that were half the acquired pixel size by zero padding in k-space. Due to the small size and slow velocity of PAs, their Gibbs ring effects are not expected to alter the phase of surrounding tissues. Therefore, CD images were calculated with Eq. (5) from the magnitude and phase difference images for convenience. The magnitude images from each coil were combined with root mean square and the phase difference images were calculated with Eq. (6). Before calculating the CD image for each PA, the phase difference and magnitude images were detrended to remove 0^th^ to 2^nd^ and 1^st^ to 2^nd^ order spatial variations, respectively, within a circular background ROI. The background ROI had a radius of 1.5 mm, centered on the PA ROI, and excluded pixels within the PA ROI or outside the WM masks. Details of PA ROI and WM mask definitions are described next.

PA ROIs were delineated with the following procedure. First, WM masks were defined on the T_1_ weighted images which were oversampled to match the pixel size and FOV of the PC images. Second, the slow spatial variations in the phase difference and magnitude images (caused by eddy current and coil sensitivity inhomogeneity) within the WM mask were estimated with a second order polynomial and then removed from the original images. Third, standard deviations (σ) of the intensities within the WM mask in the detrended phase difference and magnitude images were calculated and pixels with intensities 1.96σ above the mean were selected. Fourth, pixels above the thresholds were grouped into neighboring clusters and clusters in the phase difference images that overlapped with a cluster in the magnitude image were defined as PA ROIs. Fifth, the resulting ROIs were visually inspected to remove ROIs with irregular shapes.

*S*_*wm*_ needed for calculating *S*_*f,0*_ in Eq. (9) was calculated as the mean of the magnitude over the background ROI associated with each PA as defined above. Thus, different *S*_*wm*_ values were used for different PAs due to image inhomogeneities. T_1_, T_2_*, and γ values needed for calculating e(v) and *S*_*f,0*_ were the same as specified above for the simulation. MBAC fitting was performed by including pixels within a circular ROI of 0.6-mm radius centered at each PA. The fitting procedure was the same as in simulation.

To estimate the variability of fitted *D*, 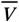, and VFR, PAs that were detected in both scans in each subject were selected and the root mean square (RMS) of the inter-scan differences (∆_r_) over all such vessels were calculated. Assuming Gaussian distributions, the RMS value of ∆_*r*_ (∆_r_,rms) is approximately 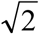 times the measurement error. Therefore, the errors of the D, 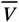, and VFR were estimated by 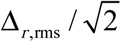.

To allow in vivo evaluation of the accuracy of MBAC-derived fitted results, the high resolution ICA images (an example in Figure 2(A)) were blurred by keeping 5% - 100% of the acquired k-space data along each spatial dimension at k-space center, resulting in image resolutions of 0.446 mm to 8.92 mm. However, signals from neighboring blood vessels can overlap with ICA in low-resolution images and affect analysis results, as shown in Figure 2(B). Therefore, a second ROI was defined to include all blood vessels near ICA as shown by the green contour in Figure 2(A), and signal within the ROI was set to zero before generating low resolution images. The reconstructed low resolution images had the same pixel size and FOV as the original high-resolution images by zero padding in k-space. Since the ICA signal is much stronger than nearby tissue, phase reversal is expected at nearby pixels. Therefore, Eq. (4) was used for calculating CD. The phase images were calculated from Eq. (13) using ICA ROI for aligning the reference image phases of different channels. The ICA ROIs were roughly circular and were manually drawn on the original high resolution magnitude images to include all pixels with clear flow induced signal enhancement compared to the neighboring tissue.

**Figure 2.**
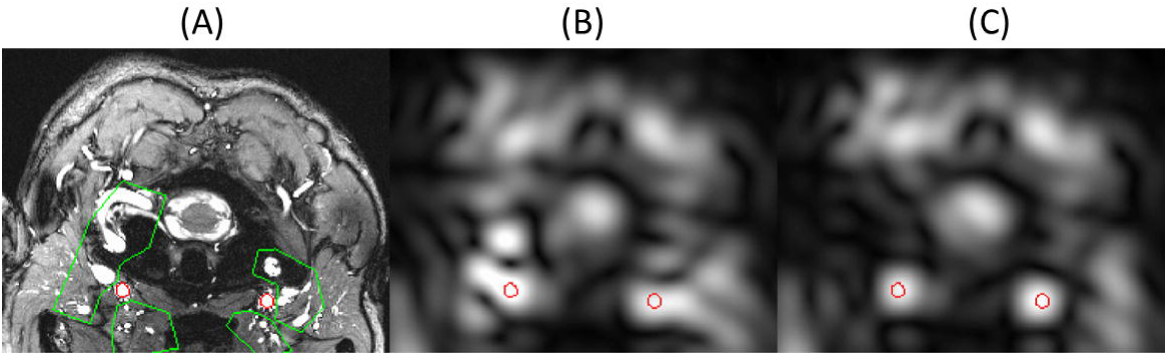
(A): High resolution magnitude image acquired perpendicular to the ICA. The ICAs are delineated by the red contours. (B) and (C): low resolution image reconstructed by keeping only the central 5% of the k-space data without and with removing the tissue signal within the green contours in (A), respectively. Without removing the neighboring vessels, their signals overlap with ICA as can be seen in (B).

MBAC was performed on pixels in an 8-mm-radius circle centered on the ICA ROI with the same fitting procedures as described above in the simulation. *S*_*f,0*_ was calculated as the mean signal intensity in the ICA ROI on the high resolution magnitude images without flow-encoding gradient divided by the theoretical flow enhancement factor calculated based on the mean velocity within the ICA ROI. The differences between MBAC-derived 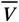, D, and VFR values and true 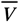, D, and VFR values were calculated and averaged over the eight ICAs in the four subjects. Since calculations of 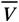 and D directly from the high resolution images are still affected by the partial volume effects of pixels at the edge of the ROIs, the true 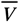, D, and VFR values were instead defined as MBAC-derived values based on the original high resolution images. The MBAC approach should also correct for the partial volume effects of the edge pixels in high-resolution images.

To evaluate whether the velocity distribution within ICA follows a laminar pattern, we separated pixels in the ICA ROIs into groups based on their distances from the pixel to the center of the ROI in the high resolution images. Then the phase differences of pixels with the same distances were averaged to study its dependence on distance to ICA center. Before averaging, the phase shift of each pixel were normalized by the mean phase shift within the corresponding ROI. The resulting phase difference versus distance curve were fitted with the theoretical expression for a laminar flow pattern 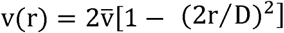 with 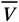 and *D* as free parameters using the lsqcurvefit function in MATLAB.

## Results

### Simulation

Figure 3 shows the velocity dependence of the noise-induced fluctuations of fitted 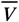, D, and VFR. The SDs increased with decreasing 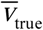 and *D*_true_, consistent with the expectation that it becomes more difficult to accurately measure the flow and vessel diameters with increased partial volume effects and decreased flow induced signal enhancement. The SD of the MBAC approach is much smaller than for the MBAP approach. At 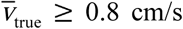 with MBAC, the SDs of 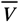, *D*, and VFR relative to their true values are ≤38%, for vessels with *D*_true_ ≥80 μm. Note that at *D*_true_ ≥ 80μm, the vessel occupies only 5.2% of the acquire pixel area.

**Figure 3.**
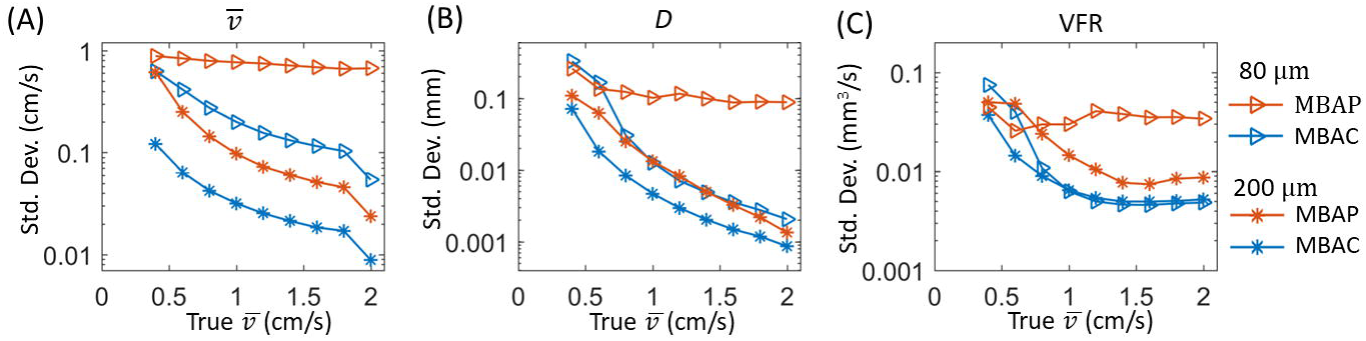
Standard deviations of fitted (A) 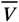, (B) *D,* and (C) VFR in the simulation as a function of true 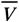, obtained with the MBAP and MBAC methods for vessel diameters of 80 and 200 um. Note that a log scale was used for the y-axes due to the large value range.

To evaluate whether there are systematic offset of the fitted values relative to the true values, the difference between mean fitted 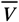, *D*, and VFR over the 100 repetitions and their true values are displayed in Figure 4. The MBAC results show no systematic offset when 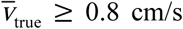. At lower 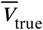, the parameters are overestimated, to a larger degree at smaller *D*_true_. In contrast, the MBAP results deviate from the true values at all 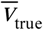 when the vessel diameter is 80 μm. When the vessel diameter is 200 μm, the deviation becomes significant only at the lowest 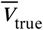 of 0.4 cm/s. The systematic errors of the fitted 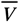, *D*, and VFR do not result from trapping of the fitting algorithm in local minimum since the residuals for the final fit were always less than the residuals for the true values, which indicate intrinsic bias of the MBAC and MBAP methods at small *D*_true_ or 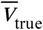.

**Figure 4.**
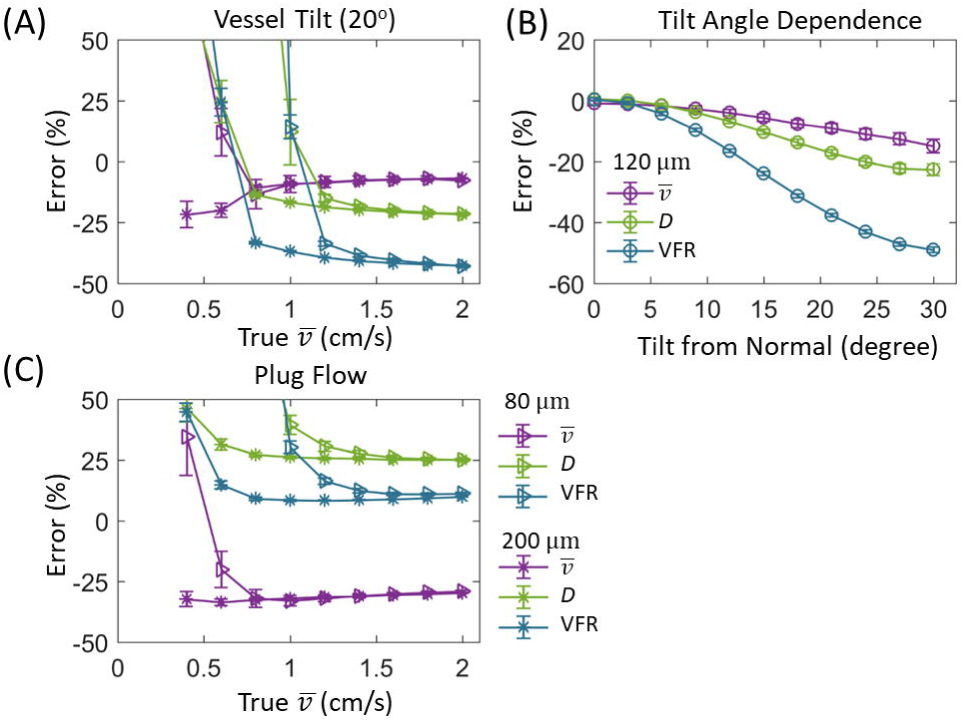
Mean difference between fitted or apparent 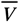 (left column), *D* (middle column), and VFR (right column) and their true values as a function of true 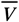. The upper and lower rows are for vessel diameters of 80 and 200 μm, respectively. The error bars are standard errors of the mean.

Figure 4 also shows the difference between the apparent and true 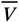, *D*, and VFR. The apparent parameters show much larger deviations from the true values compared with MBAC and MBAP results, with the deviations increasing at larger 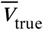, demonstrating the necessity for correcting partial volume effects. The apparent diameter and velocity show opposite changes relative to their true values. However, these two opposite changes do not cancel each other out when combined to calculate VFR, resulting in overestimation of VFR compared to the true values, especially at 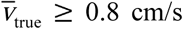.

Tilt of vessels away from the slice normal direction and deviation of flow from a laminar pattern both introduce systematic errors in the fitted results. Figure 5(A) shows the difference between MBAC-based fitted and true 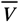, *D*, and VFR values, after normalization by the true values, as a result of 20° vessel tilt. Vessel tilt reduces the fitted 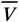, *D*, resulting in large underestimate of VFR at tilt = 20°. The normalized errors are almost independent of vessel diameter and velocity when 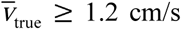, but increase at lower 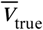, which may be related with the systematic bias as observed in Figure 4 without vessel tilt. Figure 5(B) shows the 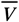, *D*, and VFR errors versus tilt angle for a vessel with *D*_true_ = 120 μm and 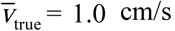, demonstrating a strong dependence of the error on tilt angle. Figure 5(C) shows the normalized error of the MBAC results when the vessel has a plug flow pattern. Similar to the tilt angle dependence, the normalized errors are almost independent of vessel diameter and velocity at large 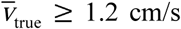, while increasing at lower 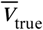. The plug flow pattern has opposite effects on fitted 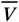 and *D*, resulting in smaller errors in VFR.

**Figure 5:**
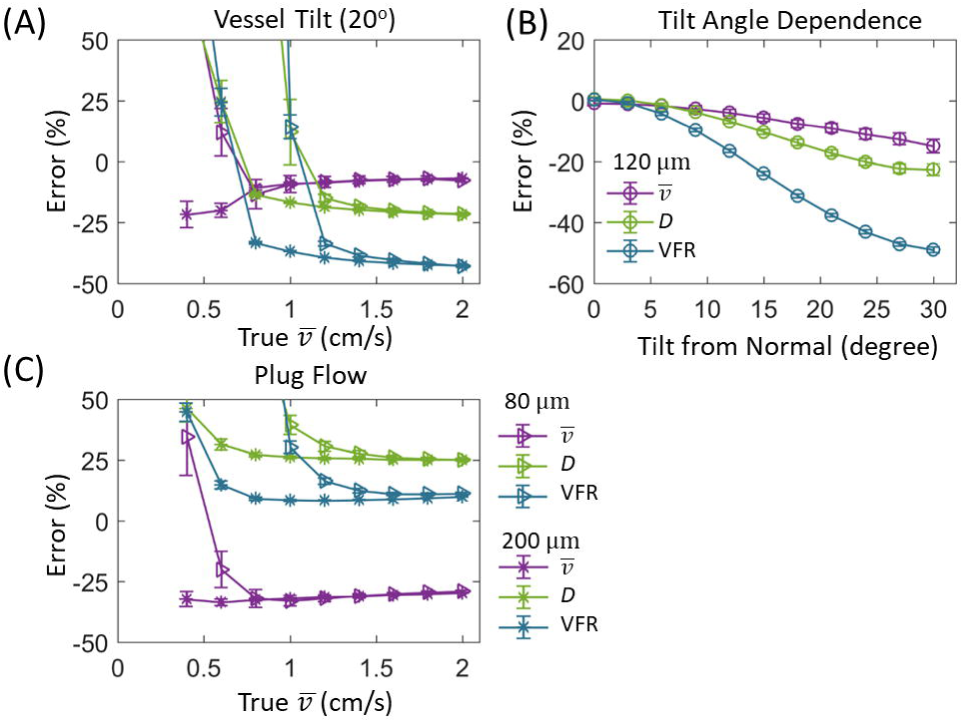
Mean difference between fitted and true 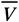, *D*, and VFR, after normalization by the true values. In (A) the underlying vessel was tilted by 20° relative to the slice normal direction and had diameters of 80 or 200 μm. In (B), the vessel had a diameter of 120 um and the differences were plotted against tilt angles. In (C), the underlying vessel was not tilted but had a plug flow profile. Different colors represent different quantities while different symbols represent different vessel diameters. The error bars are standard errors of the mean.. Note that the model images used for fitting were calculated based on perpendicular vessels with laminar flow, as is the case throughout the paper.

### Phantom Study

From the variable TR TSE sequence, a water T_1_ of 3.5 s was obtained and was used for calculating the model images in the MBAC method.

The best-fit model images from MBAC matches well with the corresponding measured images, as shown in an example in Fig. 6(A). The fitted 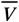, *D*, and VFR results agree well (percent difference < 51%) with the true values when 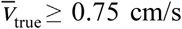 for all three flip angles, as shown in Fig. 6 (B)-(D). The errors become more variable between the three flip angles at lower 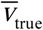, consistent with the increased fluctuations with decreasing 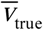 observed in the simulation. Compared to the MBAC results, the apparent 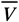, *D,* and VFR without correcting for partial volume effects show much larger errors relative to the true values, as shown by the red symbols in Fig. 6 (B)-(D).

**Figure 6:**
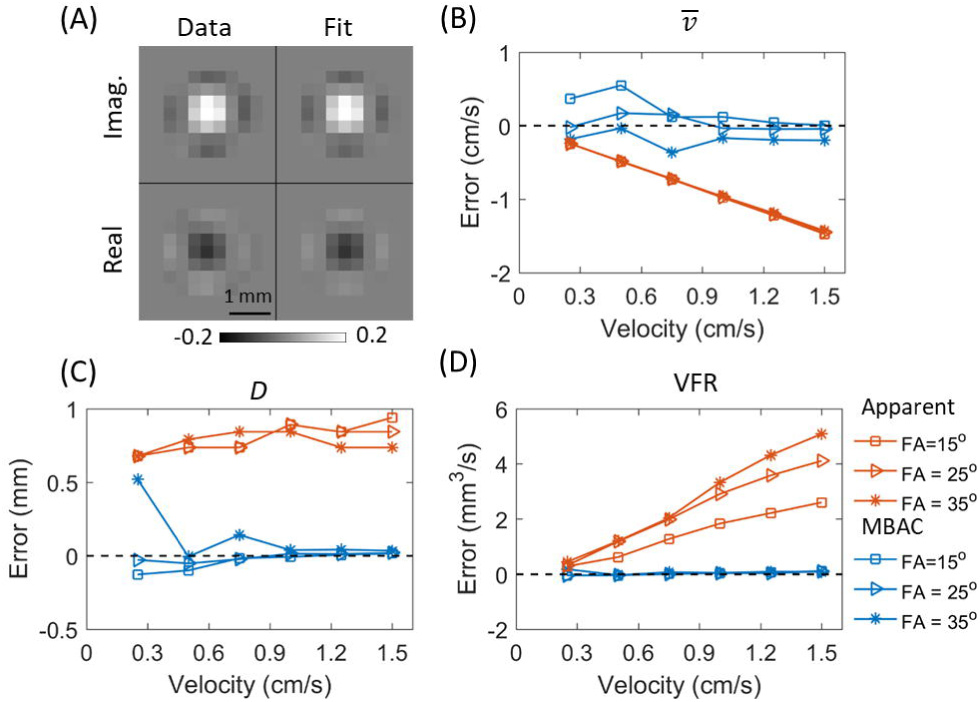
Results from the flow phantom. (A): An example of the imaginary and real parts of measured complex difference image in the flow phantom and corresponding MBAC fit. The data were acquired at a velocity of 1 cm/s and flip angle of 35°. (B) – (D): Errors between fitted or apparent 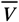, *D,* and VFR and their true values, as a function of mean flow velocity. Different symbols represent different flip angles used for data acquisition.

### In vivo Study

#### Internal Carotid Artery

The true 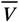, D, and VFR of the eight ICAs, defined as corresponding fitted MBAC parameter values based on the original high-resolution images, were 23.4±4.4 cm/s, 4.1±0.2 mm, and 189±40 ml/min, respectively. The fitted 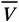, D, and VFR based on the lower-resolution images agrees well with the true values, as shown in Fig. 7(A) to (C). The mean percentage errors were ≤ 12% at all spatial resolution, down to the lowest spatial resolution (pixel size = 8.9×8.9 mm^2^), for which the lumen occupies only 18% of a pixel. The validity of the laminar flow assumption in ICA is supported by the observed velocity versus radial distance curve, which can be fitted well by the laminar distance dependence, as shown in Fig. 7(D).

**Figure 7:**
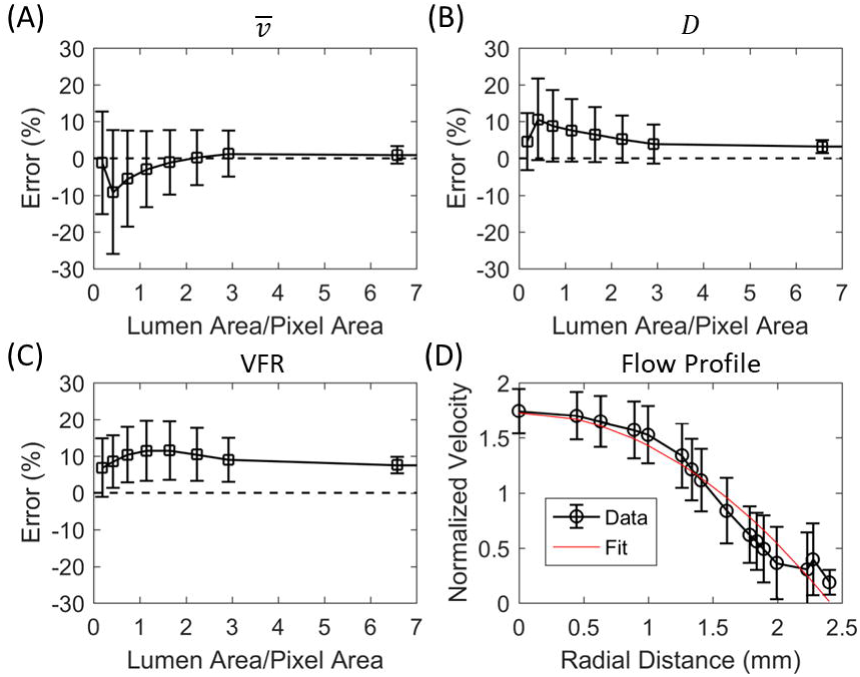
*In vivo* results from the internal carotid arteries. (A)-(C) give errors between fitted 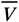, *D,* and VFR and their true values, respectively, after normalization by their true values as a function of spatial resolution. The spatial resolution is expressed as the ratio of the lumen area and the acquired pixel area. The symbols and error bars represent means and standard deviations across the 8 ICAs in 4 subjects. (D) shows the velocity of the pixels in the ICA ROIs, after normalization by the mean velocity in the ROIs, as a function of the distance between the pixels to the centers of mass of the ROIs. The results are calculated based on the original high-resolution images. Pixels at the same distance are averaged together and the error bars are standard deviations. The red line is the least square fit with the expected laminar distance dependence.

#### Penetrating Arteries in CSO

A total of 379 vessels were identified in the 12 scans, among which 252 were from 126 vessels that were identified in both scans from the same subject. Representative magnitude and phase images are displayed in Figure 8(A) to (C), where the identified penetrating arteries are enclosed by red circles. Figure 8(D) shows representative CD images for a single vessel and the MBAC fitting result, showing an excellent match between the measured and fitted images.

**Figure 8:**
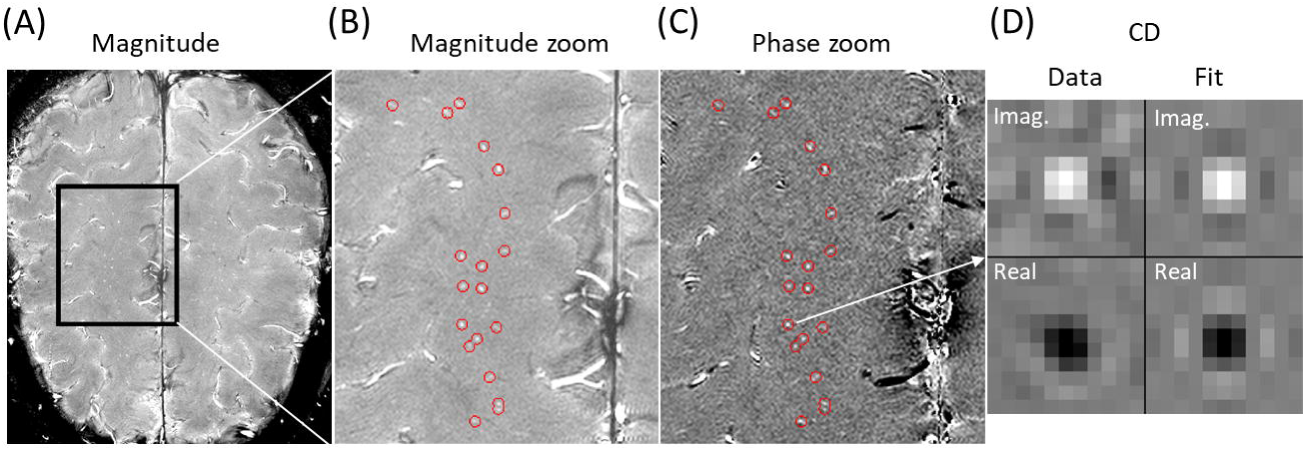
(A) shows representative magnitude image acquired with PC-MRI perpendicular to penetrating arteries in CSO. (B) and (C) are magnified magnitude and phase images, respectively, of the region enclosed by the rectangle in (A). Red circles enclose PAs identified by the thresholding method. (D): An example of the imaginary and real parts of the complex difference images (left) and corresponding fitted MBAC model images (right).

Application of MBAC on the 379 vessels gave fitted *D* distribution in the expected range (i.e. ≤500 μm (Pesce and Carli 1988)) for the majority (87%) of vessels with a minimum D of 88 μm and a peak around 120 μm. All the vessels with fitted D≥500 μm had fitted 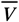 less than 0.4 cm/s. As our simulation and phantom studies suggest that MBAC results can have large uncertainties and systematic errors when 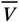 is less than 0.8 cm/s, those large diameter values are most likely artifacts of the MBAC fitting. To ensure the validity of the MBAC results, vessels with fitted 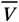 less than 0.8 cm/s were excluded from the following analysis. As a result of the exclusion criteria, 136 (36%) vessels were excluded.

Figure 9(A) – (C) display the MBAC-derived 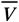, D, and VFR distributions of the remaining vessels. The 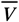 and D distributions have peaks at 1 cm/s and 0.12 mm, respectively. Most PAs have diameters between 0.1 mm and 0.2 mm and velocity below 2.5 cm/s. As a comparison, the distributions of apparent 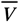, D, and VFR without correction of partial volume effects of the same vessels are shown in Figure 9(D) – (F). Consistent with the simulation and phantom results, the velocities are greatly underestimated by the apparent values, while the diameters and VFRs are greatly overestimated. The means and standard deviations of the distributions are given on the first data row of Table 1.

**Table 1.** The mean and standard deviations of the 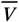, D, and VFR distributions and measurement errors of 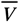, D, and VFR for single vessels (rows 1 −3) and for their mean values after averaging within subjects.

|  |  | MBAC |  |  | Apparent |  |  |
| --- | --- | --- | --- | --- | --- | --- | --- |
| | | $\bar{v}$ (cm/s) | D (mm) | VFR (mm<sup>3</sup>/s) | $\bar{v}$ (cm/s) | D (mm) | VFR (mm<sup>3</sup>/s) |
| Vessel Level | Mean (STD) | 1.3 (0.4) | 0.14(0.03) | 0.20(0.09) | 0.34 (0.10) | 0.54(0.11) | 0.87(0.59) |
|  | Error | 0.28 | 0.02 | 0.024 | 0.045 | 0.06 | 0.23 |
|  | Error/Mean | 22% | 14% | 12% | 13% | 11% | 26% |
| Subject Level | Mean (STD) | 1.3 (0.14) | 0.14(0.02) | 0.21(0.05) | 0.36(0.09) | 0.55(0.05) | 1.0(0.49) |
|  | Error | 0.16 | 0.017 | 0.031 | 0.029 | 0.041 | 0.32 |
|  | Error/Mean | 12% | 12% | 15% | 8% | 7% | 32% |

**Figure 9:**
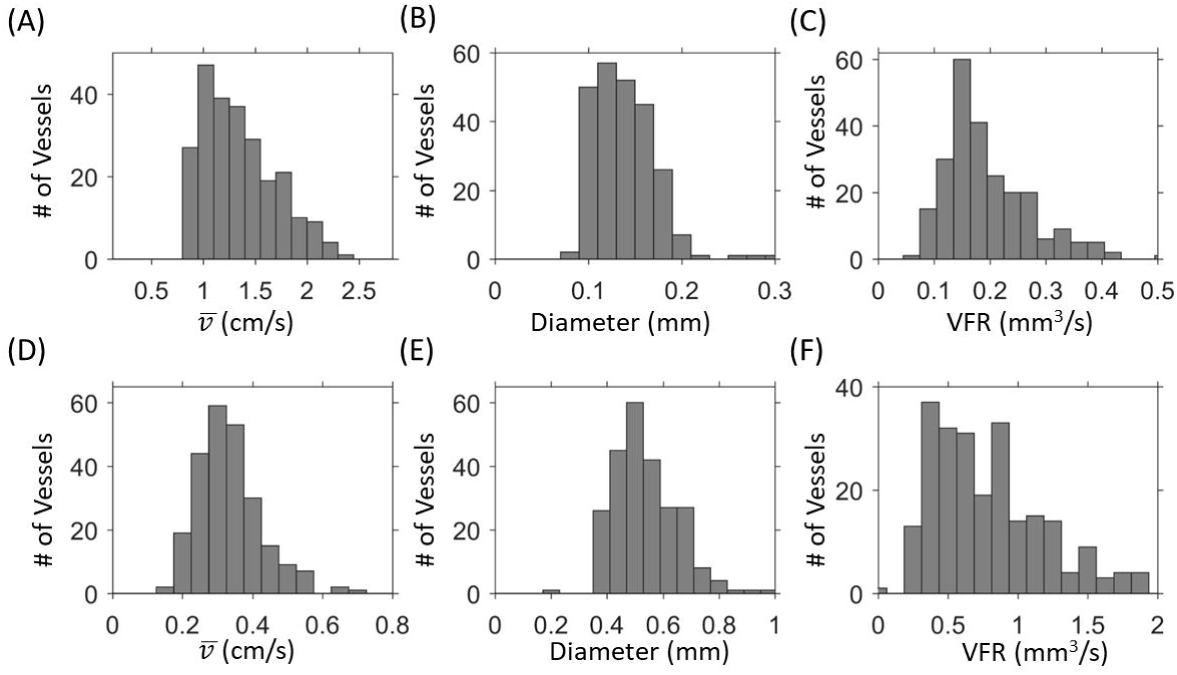
MBAC-derived (A – C) and apparent 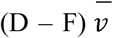, *D*, and VFR distributions of all the vessels identified in the 6 subjects. Note the differences in data ranges between the upper and lower panels. Only vessels with MBAC fitted 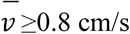 are included.

The errors of the fitted and apparent 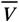, *D*, and VFR in individual vessels were calculated based on inter-scan differences of vessels identified in both scans. After excluding vessels with fitted 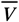 less than 0.8 cm/s in either of the two scans, 72 of the 126 vessels remained for error calculation. The estimated errors for 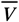, *D*, and VFR are given on the second data row of Table 1. Note that the errors of the MBAC results are much larger than the corresponding errors obtained from the simulation for a vessel with similar 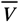 and *D.* Based on the simulation, a vessel with *D*_true_ = 0.14 mm and 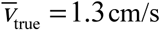 has errors for 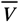, *D*, and VFR of 0.04 cm/s, 0.004 mm, and 0.006 mm^3^/s, respectively. To compare the errors between the fitted and apparent parameters, the ratios of the errors to their mean values are given in the third data row of Table 1. The relative errors are comparable between the MBAC and apparent values, suggesting that the MBAC method does not lead to amplification of parameter errors.

In addition to vessel level analysis, the mean 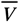, *D*, and VFR (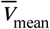, *D*_mean_, and VRF_mean_) over all vessels (after excluding those with 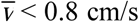) in a scan can be calculated and may serve as a useful index for PA characterization for each subject. The group averaged 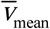, *D*_mean_, and VRF_mean_ over the 12 scans are given in the fourth row of Table 1. The errors of 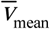, *D*_mean_, and VRF_mean_ based on the inter-scan variations within each subject are given in the fifth row of Table 1. The results suggest that the vessel-averaged MBAC results have relative errors ≤ 15%, smaller than errors for single vessel results.

## Discussion

In this paper, we carried out simulation, phantom, and in vivo studies to evaluate the proposed MBAC method for estimating 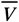, D, and VFR from phase contrast MRI. According to our simulation results, our method has increased precision and accuracy compared with an earlier method utilizing only phase images,(Hoogeveen, Bakker et al. 1999) enabling quantitative measurement of vessels with severe partial volume effects. As long as the velocity is ≥0.8 cm/s, relative errors ≤31% can be achieved for vessels occupying only 5.2% of imaging pixel area, much smaller than achieved earlier with the MBAP method. (Hoogeveen, Bakker et al. 1999) The improvements of MBAC method over MBAP can be explained by the doubling of the data points for the model fitting in MBAC. It is important to note that in order to realize the doubling of data points, the PC MRI sequence should acquire two images with the velocity encoding gradient on and off, instead of acquiring two images with opposite polarity of the gradients. In the latter case, the real part of the CD image is always zero. Furthermore, since static tissue signals are absent in CD images, our method does not entail an appropriate modeling of the static tissues and is therefore more suitable for PAs where signals from nearby vessel wall, perivascular spaces, and white matter can overlap with PAs due to the limited spatial resolution.

We further evaluated the accuracy and precision of our approach with phantom and in vivo studies. The flow phantom and ICA results demonstrate that the MBAC method can get accurate estimation of 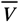, *D*, and VFR in presence of strong partial volume effects. In the flow phantoms whose cross sectional area is only 17% of the acquired pixel area, the discrepancies between the measured and true 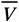, *D*, and VFR values were less than 51% when flow velocity is ≥ 0.75 cm/s which is the velocity range for the majority of PAs detected in our study. In the ICA, the discrepancies were less than 12% with a similar lumen to pixel area ratio of 18%. The precision of the MBAC method for measuring PAs was evaluated by inter-scan variations of 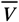, D, and VFR. Our results suggest that relative errors of 22% or less can be achieved for PAs with 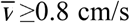, where the PAs on average occupy 18% of the acquired pixel size. The errors are higher than estimated from the simulation, which is expected since the simulation only considered errors caused by random thermal noise but not other scan to scan variations such as different motion artifacts between scans which may lead to artificial vessel diameter increase due to blurring and/or reduced 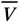values due to reduced phase contrast in the image. In the future, prospective motion correction techniques based on fat navigator echoes may be incorporated into the sequence to reduce motion artifacts.(Gallichan, Marques et al. 2016)

Our simulation, phantom, and in vivo PA results all demonstrate that MBAC method has large errors for vessels with velocity <0.8 cm/s. This can be explained by the poor signal to noise ratio at low velocities, resulting from decreased flow enhancement effect and phase shifts. Given low SNR, the MBAC method become unable to converge to the true parameter values and the fitted D values not only have large uncertainties but also contains a systematic error which results in overestimate of *D*. The overestimation of *D* at low 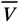 are also consistent with the in vivo results where all the vessels with fitted *D*≥500 μm had fitted 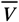 less than 0.4 cm/s. Therefore, to ensure the accuracy of the MBAC method for PA measurements, vessels with fitted 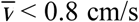 should be excluded in future studies.

Using the MBAC approach, the 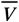, D, and VFR distributions of PAs were obtained for the first time in young healthy adults, which differ greatly from results without correcting for partial volume effects as shown in Figs. 9, consistent with simulation and phantom results in Figs. 4 and 6. The mean 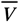 is about twice the values reported earlier without correction for partial volume effects.(Geurts, Biessels et al. 2018) As expected, contamination from static tissue reduces the phase shift induced by the velocity encoding gradients, resulting in a lower apparent velocity. Most PAs in our study have diameters in the range of 88 – 200 μm, consistent with earlier ex vivo studies.(Fisher 1968, Pesce and Carli 1988) Fisher observed PAs in the basal ganglia and pon with diameters in the ranges between 40 – 200 μm,(Fisher 1968) while Pesce and Carli reported a diameter range of 100 – 536 μm in the putamen with the majority of PA diameters between 100 – 200 μm.(Pesce and Carli 1988)

As the partial volume effects depend on signal contamination from nearby tissues, which might be altered during SVD pathogenesis, the apparent 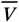, D, and VFR are unsuitable for monitoring the PA abnormalities. In contrast, the MBAC method removes nearby tissue signal contaminations and allows true 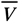, D, and VFR to be measured, which may serve as useful indices for investigating the role of pathological PA alterations in the development of cerebral SVD. For example, if there exists hypoperfusion in the deep WM caused by narrowing of the PA lumen, a reduction of PA diameter and VFR are expected prior to the appearance of typical WM changes in MRI such as lacunar infarct and hyperintensity in T_2_-weighted images.(Yao, Sadoshima et al. 1992, Bernbaum, Menon et al. 2015) On the other hand, no early diameter and VFR reduction should be observed if leakage of blood-brain barrier due to endothelial failure is the underlying etiology of SVD.(Wardlaw, Sandercock et al. 2003)

There are several limitations of the current study. First, all PAs were assumed to be perpendicular to the imaging plane. According to our simulation, this assumption can lead to an underestimation of 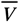, *D*, and VFR for tilted vessels. The orientation of PAs may be inferred from the orientation of the surrounding PVS which can be visualized with a high resolution 3D T_2_ weighted MRI.(Bouvy, Biessels et al. 2014, Zong, Park et al. 2016) The measured angle of PAs can then incorporated into the MBAC image model for deriving more accurate 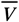, *D*, and VFR values. Second, *D* and VFR might decrease along the PA path as blood flow is diverted into smaller arteriole braches. In the future, the slice-position dependence of *D* and VFR should be characterized. Third, due to limited signal to noise ratio, not all PAs can be visualized and PAs with 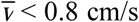 cannot be measured accurately and have to be excluded. Therefore, the histograms in Figure 9 are likely underestimated at small 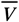, D, and VFR values. Fourth, non-laminar flow pattern can lead to underestimate of flow velocity and overestimate of vessel diameter, according to our simulation. In the extreme case of completely uniform velocity, the error is ~32%. However, because of the small Reynold number (*R*_*e*_) and short entrance length (~0.05*DR*_*e*_ (Bergman and Incropera 2011)) where non-laminar flow is present, the deviation from laminar flow at the slice location and thus the associated fitting error might be negligible.

## Conclusion

We have developed a new MBAC approach for quantitative measurement of velocity, diameter, and VFR of penetrating arteries in CSO. Our simulation results demonstrate much smaller measurement errors of MBAC compared with model based analysis of phase images. The technique has been validated in a flow phantom and in vivo on internal carotid arteries. Measurement errors and distributions of velocity, diameter, and VFR were obtained in PAs. Our study suggest that PAs with velocities ≥ 0.8 cm/s can be quantitatively measured in the presence of severe partial volume effects with the MBAC method, which may serve as an invaluable tool for investigating the role of PAs in the aetiopathogenesis of cerebral small vessel disease.

## Acknowledgements

This study was supported by NIH grant 5R21NS095027-02. We thank the BRIC staff members for their support.

